# The generalized spatial representation in the prefrontal cortex is inherited from the hippocampus

**DOI:** 10.1101/2021.09.30.462269

**Authors:** Michele Nardin, Karola Kaefer, Jozsef Csicsvari

**Author notes:** Donders Institute for Brain, Cognition and Behaviour, Radboud University, Postbus 9010, Nijmegen 10 6500GL, Netherlands (Karola Kaefer).

## Abstract

Hippocampal and neocortical neural activity is modulated by the position of the individual in space. While hippocampal neurons provide the basis for a spatial map, prefrontal cortical neurons generalize over environmental features. Whether these generalized representations result from a bidirectional interaction with, or are mainly derived from hippocampal spatial representations is not known. By examining simultaneously recorded hippocampal and medial prefrontal neurons, we observed that prefrontal spatial representations show a delayed coherence with hippocampal ones. We also identified subpopulations of cells in the hippocampus and medial prefrontal cortex that formed functional cross-area couplings; these resembled the optimal connections predicted by a probabilistic model of spatial information transfer and generalization. Moreover, cross-area couplings were strongest and had the shortest delay preceding spatial decision-making. Our results suggest that generalized spatial coding in the medial prefrontal cortex is inherited from spatial representations in the hippocampus, and that the routing of information can change dynamically with behavioral demands.

## Introduction

The question of how space is encoded by networks of interacting neurons is arguably one of the most actively studied subjects in neuroscience. One of the clearest neural correlates of space is the spatially-tuned firing of hippocampal place cells. The activity of these cells provides a neural representation of the current position within an environment that is required for spatial navigation (1, 2). Although the most spatially-tuned cells reside in the hippocampus, other brain areas have also been shown to exhibit place-related firing (3–8), highlighting that the neural representation of space is distributed throughout the brain.

One of these areas is the medial prefrontal cortex, where a significant fraction of neurons also show spatial selectivity (9, 10) and play a role in spatial navigation (11, 12). The medial prefrontal cortex receives direct projections from the hippocampus, most prominently from its ventral subregion (13). Whilst it does not project back directly, the medial prefrontal cortex has indirect and bidirectional connections with the hippocampus via the thalamic nucleus reuniens (14–16) and the perirhinal and lateral entorhinal cortices (13, 17).

Given the bidirectional connection between the hippocampus and medial prefrontal cortex, a question which arises is whether the spatial information in the medial prefrontal cortex is inherited from the hippocampus. Indeed, selectively inactivating the direct projections from the hippocampus leads to deficits in the spatial coding of medial prefrontal neurons and in the performance in a spatial working memory task (18). Even more globally, encoding of spatial information in several neocortical regions seems to depend on an intact hippocampus (19). Alternatively, the spatial representations in the hippocampus and medial prefrontal cortex may emerge from bidirectional interactions (12, 20). The fact that their spatial codes are coherent at a theta cycle timescale (21) supports this possibility. Furthermore, the medial prefrontal cortex influences hippocampal coding, since inactivating the medial prefrontal cortex or the indirect nucleus reuniens projections leads to a decrease in hippocampal place cell firing variability (22), rule-based object selectivity (23) and trajectory-dependent firing (11). Until now, however, it is not clear how spatial representations in prefrontal cortical areas are established.

Previous reports characterized the spatial information conveyed by medial prefrontal neurons, and showed that both single neurons and population activity provide a generalized spatial code (24–26). Here, we tested whether generalized representations in the medial prefrontal cortex result from a bidirectional interaction with, or are mainly derived from spatial representations already present in the hippocampus. We therefore examined simultaneously recorded hippocampal and prefrontal neurons to investigate how generalized spatial representations in either region predicted its later emergence in the other. We found prefrontal cortical representations to be coherent, after a short delay, with those of the hippocampus. Moreover, by employing rigorous statistical inference methods (27), we identified pairs of cells in the hippocampus and medial prefrontal cortex that formed statistically significant and stimulus-independent cross-area couplings. The majority of such cross-area pairs involved prefrontal units lagging behind hippocampal units. Moreover, cross-area couplings inferred from data resembled the predicted optimal connections of a simple probabilistic model of spatial information transfer and generalization. Finally, cross-area couplings were strongest and shortest in delay upon spatial decision-making. Our results suggest that generalized spatial coding in the medial prefrontal cortex is inherited from spatial representations in the hippocampus, and that the routing of information can change dynamically with behavioral demands.

## Results

### Rule switching task

We analyzed the activity of simultaneously recorded single neurons from the prelimbic area of the medial prefrontal cortex (mPFC) and the CA1 of the dorsal hippocampus of four rats across 13 experimental sessions (25). Rats were trained on a rule-switching task on a plus maze where reward had to be collected following a spatial- or light-guided strategy (Fig. 1A). At the beginning of every trial, the animal was placed in one of the two start arms (north or south) to then approach the maze center and collect a reward in one of the two goal arms (east or west). Every recording day started with a block of trials with the previous day’s last rule (“old rule”) in play. This was followed by 40 minutes of sleep/rest. The next block of trials initially started off with the old rule followed by an unexpected switch in the rules (“rule switch”). The animal then had to abandon the old rule and, through trial and error, switch strategies in order to successfully perform trials according to the “new rule”. After another sleep/rest period rewards had to be collected following the new rule (Figure 1B). The point of successful rule switch was defined from the third trial of five consecutively correct trials performed according to the new rule. Animals typically took an average of 30 trials (range 10 to 77) to successfully adapt to the new rule.

**Fig. 1.**
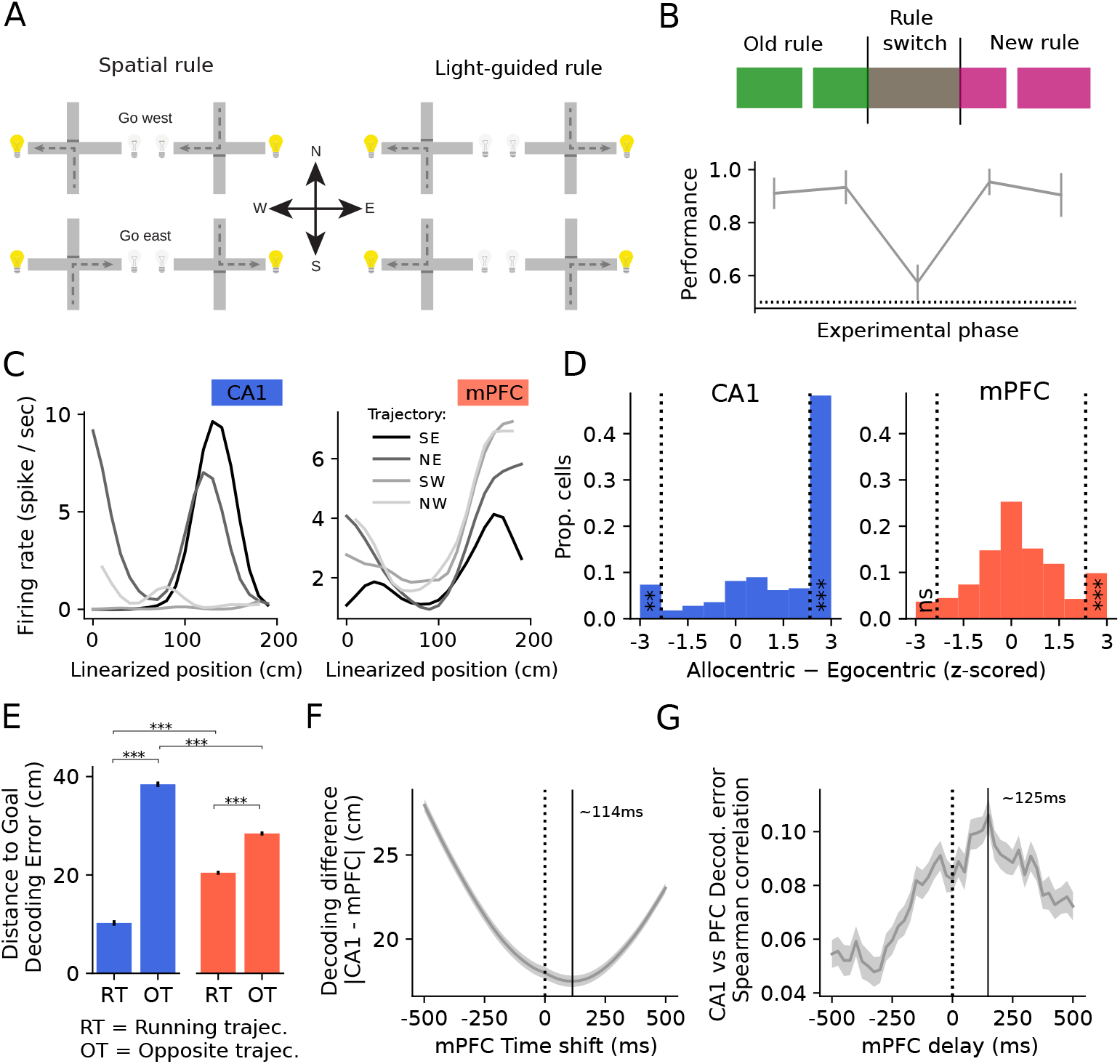
Generalized spatial information in the mPFC is influenced by that of the hippocampus. **(A)** Schematic of the behavioral rules. **(B)** Behavioral performance across sessions (N=13) for each experimental epoch. Error bars represent the 95th confidence interval for the mean across 13 sessions. Dotted line represents chance performance (p=0.5). **(C)** Example of trajectory dependent linearized firing rate maps for one CA1 (left, blue) and one mPFC (right, red) pyramidal cells. Each line represents the average firing as a function of the linearized position on the maze for one of the 4 trajectories. **(D)** Distribution of z-scored allocentric minus egocentric scores for each single cell in CA1 (left) and mPFC (right). Vertical dotted bars denote significance threshold. All significant allocentric cells are grouped in the right-most bar, all the significant egocentric cells in the left-most bar. CA1: allocentric 50.07%, Binomial test p<0.001; egocentric 7.8% (Binomial test p=0.00108). mPFC: allocentric 10.2% (Binomial test p<0.001); egocentric 4.4% (Binomial test p=0.673). **(E)** Decoding distance to goal performance of the two areas (left: CA1, right: mPFC) when using the correct trajectory (RT) or the opposite (OT). All comparisons: Mann–Whitney U test p<0.001. **(F)** Alignment of spatial representation between the two areas as a function of mPFC time shift. **(G)** Spearman correlation of decoding errors of the two areas as a function of mPFC response delay. In panels **F)**, **G)**, the shaded area corresponds to the 95th percentile confidence interval of the mean over 13 sessions. Throughout the figure, ns = not significant, ** = p<0.01, *** = p<0.001.

### Characterization of mPFC generalized spatial representation

We have previously reported (25) that mPFC cells can exhibit symmetric place fields, where cells show similar firing patterns on opposite arms of the plus maze (Fig. 1C). Here we analyzed the spatial properties of mPFC cells further. First, we explored the possibility that symmetric firing in mPFC could arise from an egocentric code. In an egocentric reference frame a cell jointly codes for position along the arm and the turn, instead of a standard allocentric position. In this case, one would observe the same firing pattern on opposite goal arms for the same turn and different firing patterns on the same goal arm on different turns. We quantified this hypothesis by measuring the similarity of the two firing rate maps of the same goal arm using data from trials where the animal took different turns (allocentric score) and subtracted the similarity of the firing rate maps of opposite goal arms using data from trials where the animal took the same turn (egocentric score). We tested the real score against the same score calculated from 1000 rate maps where trial identities were randomly shuffled. We defined cells to be significantly allocentric if their score was higher than the 95th percentile and egocentric if their score was lower than the 5th percentile of their respective shuffled score distribution. We found that mPFC cells did not show significant egocentric coding, and only a minority showed significant allocentricity (Fig. 1D), again corroborating the predominant generalization of mPFC cells (25). Surprisingly, we found that a small, yet significant proportion of CA1 cells showed egocentricity and as expected the large majority presented an allocentric coding (Fig. 1D). Next, we quantified the trajectory specificity of the hippocampal and prefrontal spatial code. We began by asking whether decoding the distance to the goal improved by using a trajectory-dependent distance-to-goal bayesian decoder, instead of a trajectory-independent one (Supp. Fig. 1A-B; methods). In CA1, using a trajectory-dependent decoder improved decoding quality as compared to a trajectory-independent one, whereas doing the same for the mPFC decreased the decoding performance (Supp. Fig. 1C). Afterwards, we asked how exchangeable the firing rate maps were across trajectory. We thus trained a decoder on one trajectory only (for example, south to east), and decoded runs from the opposite trajectory (for example, north to west). This greatly reduced the accuracy of decoding from CA1 activity (ratio 0.263), whereas it reduced only marginally the performance when decoding from mPFC activity (ratio 0.712) (Fig. 1E). These results confirm and expand on our knowledge of mPFC spatial representation, which is mostly trajectory independent and more abstract than CA1.

### mPFC’s generalized spatial representation lags behind CA1’s

Next, we explored the possibility that the spatial representation in the mPFC is inherited and transformed from CA1. In that case one should observe a delay in the mPFC representation as compared to the CA1 representation, and a correlation in the errors between the two areas (21). We decoded the distance to goal from the activity of CA1 and mPFC cells and measured the average distance between decoded positions from the two areas (Fig. 1F). The position decoded from mPFC lagged behind that of CA1, and the best alignment was observed with a mPFC delay of ~ 114ms. We then measured the correlation of decoding errors from the two populations as a function of mPFC delay. We found the best correlation at a mPFC response delay of ~125 ms (Fig. 1G), suggesting that the generalized mPFC spatial representation might be derived from CA1. To further corroborate this idea, we employed a transfer entropy measure among CA1-mPFC cell pairs (28). Transfer entropy is a non-parametric measure of the amount of directed (time-asymmetric) transfer of information between two processes. We applied this measure on the binarized spike trains of each CA1-mPFC cell pair, and found the transfer entropy to be higher in the direction CA1→mPFC than in the mPFC→CA1 direction (Supp. Fig. 1D). This supports the notion of directional information flow from the CA1 to the mPFC.

### Cross-area coupling between pairs of CA1 - mPFC cells

The analyses discussed in the previous paragraph suggested that mPFC spatial representations were derived from hippocampal ones. It is possible that not all mPFC cells are equally influenced by hippocampal activity, and similarly that not all hippocampal cells may have a strong influence on mPFC activity. To detect possible functional connections between cells across the two regions we tested whether cells exhibited correlated activity independent of common sensory inputs. This measure is typically referred to as “noise correlation”, and quantifies the amount of pairwise correlation that is not explained by similar stimulus selectivity (29). We employed a statistical estimator of noise correlations similar to the one previously introduced in (27). For each cell we fitted a generalized linear model (GLM) (30, 31) that included all possible neural and behavioral covariates that could influence and explain cross-area correlations. Those covariates were: spatial position and trajectory, theta selectivity, speed selectivity, spiking history, and within-area spiking of other cells (i.e. mPFC cells were fitted with the spiking of other mPFC cells only and, separately, CA1 cells with the spiking of the other CA1 cells only; for further details see Methods). The models were used to simulate the activity of each cell 10000 times and, for each CA1-mPFC cell pair, a cross-correlogram of the responses was computed (Fig. 2A). These surrogate cross-correlograms were used to measure by how much the actual pairwise correlation measured on real data differed from the correlation expected from simple tuning similarity (Fig. 2B). We employed this (normalized) difference as a measure of noise correlation (27). Doing this for each possible mPFC delay in the range of ± 1 sec we found that the highest number of significant pairs (i.e. noise correlation outside of their own confidence interval) was obtained when mPFC was delayed by ~50ms (Fig. 2C). Next, we characterized cells exhibiting significant noise correlations. Cells in one area that showed a significant noise correlation to at least one cell in the other area, with a maximum delay of ±200*ms*, were termed “cross-coupled”. We found that cross-coupled mPFC cells were better at decoding spatial position compared to non-coupled ones (Supp. Fig. 2A-B). In addition, the average mutual information between single cells’ activity and the distance to the goal was larger for the cross-coupled cells than that for the non-coupled cells in mPFC (Fig. 2D). Cells in CA1 that were cross-coupled were better at decoding the relative position between the start and goal, whereas non-coupled CA1 cells were better at discriminating the 2D position in the maze (Supp. Fig. 2A-B,E). These findings indicate that mPFC cells are preferentially coupled with CA1 cells that generalize spatial information across trajectories. We therefore measured, separately for the two subpopulations (cross-coupled vs non-coupled), the similarity of their spatially dependent firing on opposite arms, and found that cross-coupled CA1 cells were more symmetric than their non-coupled peers (Fig. 2D); mPFC symmetry of cross-coupled cells did not differ significantly from non-coupled ones (Supp. Fig. 2D). We explored the potential mechanisms by which neurons in mPFC might be cross-coupled. It is well-known that CA1 and mPFC undergo theta oscillation synchrony (32); we therefore asked whether mPFC cross-coupled cells showed a higher level of spiking selectivity to hippocampal theta oscillations. We found a positive relation between absolute coupling strength and hippocampal theta selectivity in mPFC, but not in CA1 (Supp. Fig. 2C).

**Fig. 2.**
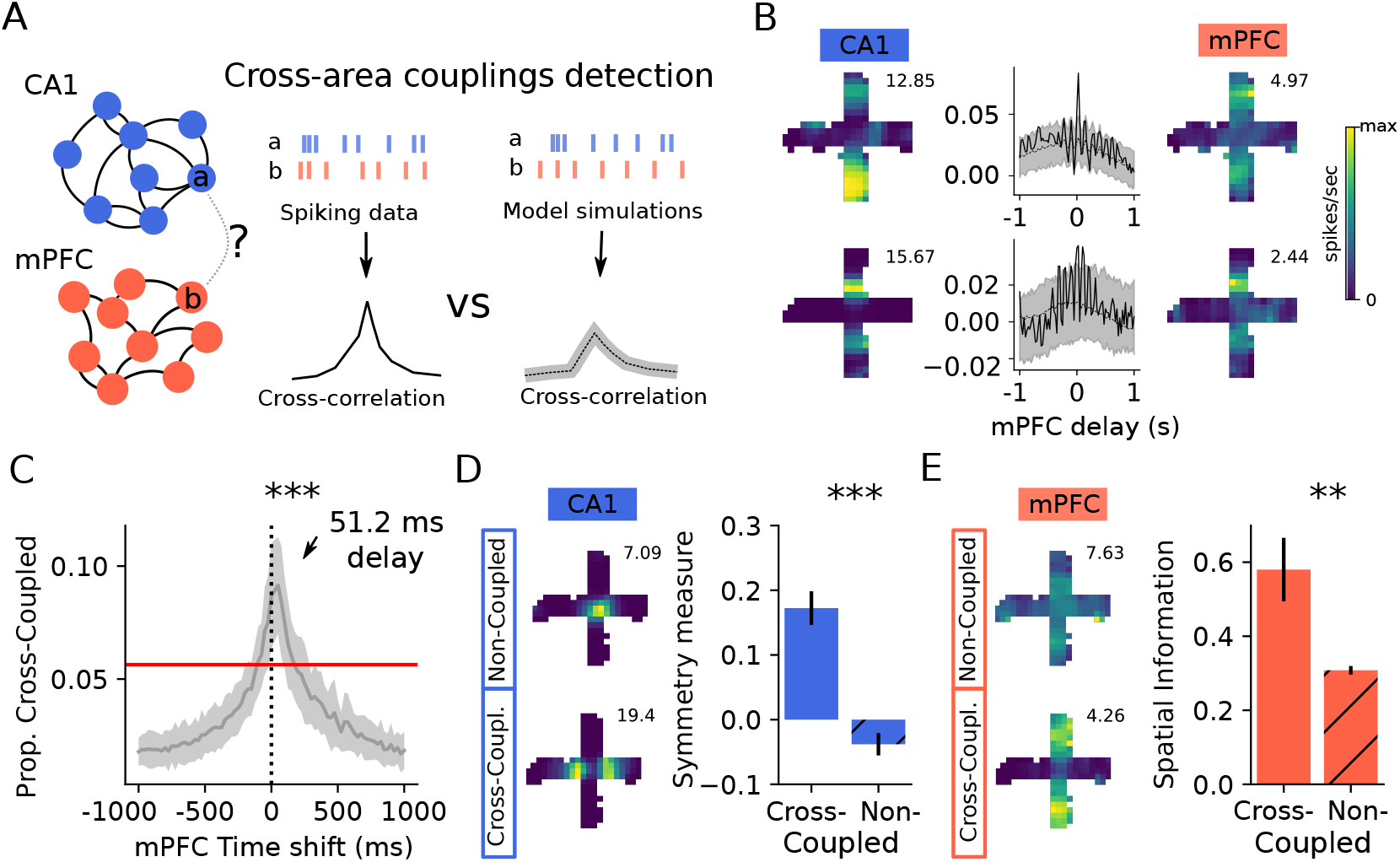
Cross-area couplings between CA1 - mPFC cell pairs. **(A)** Schematics for the detection of cross-area couplings. We employed a GLM null model that captured trajectory dependent firing, theta and speed selectivity, within-area couplings and spiking history. GLM null models were used to simulate firing 10000 times, and create a null-distribution over cross-correlograms. **(B)** Examples of cell-cell significant couplings across areas. Left and right: CA1 and mPFC 2D firing rate map. Center: cell-pair cross correlogram computed on real data (solid black line) vs shaded area representing amount of cross-correlation explained by the null model (dotted black: average, shaded area: ± 3STD). **(C)** Proportion of significant couplings as a function of mPFC delay. A pair was deemed significant at a given delay t if the real data cross-correlation at t exceeded +3STD or fell below -3STD of the average cross-correlations at t computed on simulated data. Red line represents the average+5STD of significant couplings over all possible cell pairs and delays >500ms. Shaded area represents the 95th confidence interval for the mean proportion of cross-coupled cells computed across 13 sessions. **(D)** Left: example of non- (above) and cross- (below) coupled cells in CA1. Right: CA1 PV symmetry measure of cross-coupled vs non-coupled subpopulations. Error bars represent the 95th confidence interval for the mean over all PVs across locations and sessions. Mann–Whitney U test p<0.001. **(E)** Left: example of non- (above) and cross- (below) coupled cells in mPFC. Right: average single cell spatial information of mPFC cells (right). Error bars represent the 95th confidence interval for the mean over all mPFC cells. Mann–Whitney U test p<0.01. Throughout the figure, ns ** = p<0.01, *** = p<0.001.

### Potential mechanisms of spatial information transfer and generalization from CA1 to mPFC

Next, we employed a statistical modeling approach to gain insight into a possible mechanism through which mPFC spatial activity might be derived from hippocampal representations through varying degrees of functional connections. We employed a model of information transfer and generalization; the model consisted of two layers of units with feedforward and internal connectivity, so that the second layer’s output is a function of the input received from the first layer and internal dynamics. We modeled the units in the first layer to be selective to both spatial location and trajectory (“*selective* units”, i.e., idealized CA1 place cells), whereas units in the second layer possessed fixed internal connectivity but no ab initio spatial selectivity (“*abstract* units”, i.e., idealized mPFC cells) (Fig. 3A). Our goal was to find connections from *selective* to *abstract* units such that the latter units could represent a trajectory-independent, hence abstract, spatial position. This model is probabilistic and minimally structured (i.e. maximum entropy; (33)), and resembles a Boltzmann machine in its formalization (34). The aim of this simplistic model is not to be biophysically realistic, but rather to utilize it as a statistical tool to create predictions and validate our observations.

**Fig. 3.**
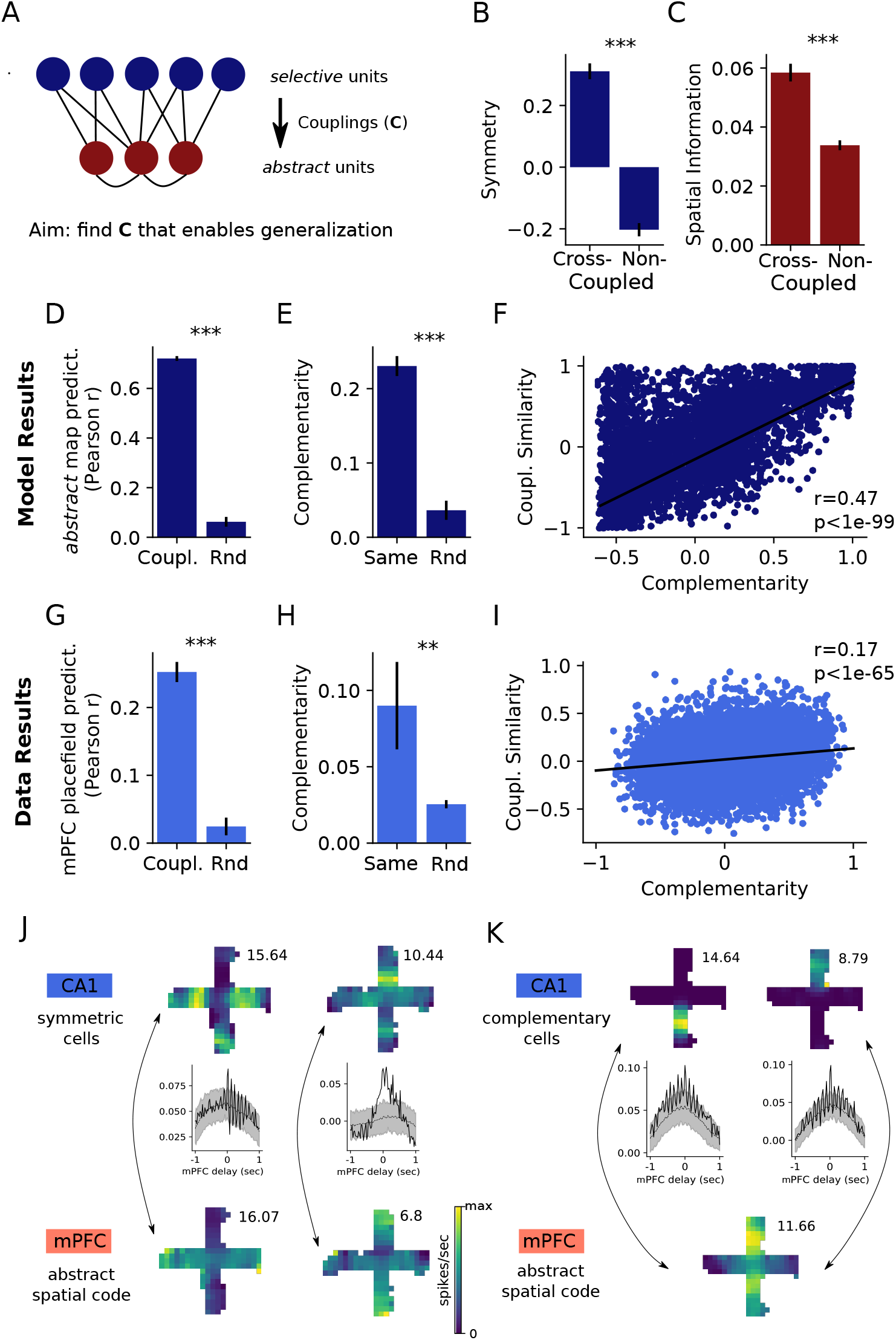
Cross-area couplings enable generalization. **(A)** Schematics of information transfer and generalization stochastic model. The input layer is composed of *selective* units (i.e., CA1-like) exhibiting trajectory dependent spatial firing. The second layer is composed of *abstract* (i.e., mPFC-like) units, whose activity is influenced by the first layer and internal connectivity. We found the matrix **C** by maximizing the mutual information between the *abstract* population firing and distance to the goal. **(B)** *Selective* units with strong cross-couplings had higher symmetry (i.e., similar firing across trajectories). Here and in the following, error bars represent the 95th confidence interval for the mean. Mann–Whitney U test, p<0.001. **(C)** *Abstract* units with strong cross-couplings exhibited higher spatial information. Mann–Whitney U test, p<0.001. **(D)** Prediction of *abstract* units firing map by using *selective* units firing maps weighted by optimized coupling matrix (left) or random couplings (right). Mann–Whitney U test, p<0.001. **(E)** Complementarity of the two *selective* units with strongest coupling to the same *abstract* unit (left) vs random (right). Mann–Whitney U test, p<0.001. **(F)** Complementarity of all *selective* pair vs coupling similarity to *abstract* units. Pearson correlation test, p<0.001, r=0.47. **(G)** Same as **D)**, for real data mPFC place fields. Mann–Whitney U test, p<0.001. **(H)** Same ad **E)**, for real data cross-couplings. Mann–Whitney U test, p<0.01. **(I)** Same as **F)**, for real data CA1 cell pairs. Pearson correlation test, p<0.001, r=0.17. **(J)** Generalization mechanism 1. CA1 cells with symmetric firing fields (top row, two example firing rate maps, peak firing rate in topright corner) show significant cross-couplings with mPFC cells (bottom row, two examples). Middle row shows the cross correlograms between top and bottom cells. (black line = real cross-correlation, black-dotted = average crosscorrelation from null-model simulations, shaded ares = ±3STD of the mean from null-model simulations). **(K)** Generalization mechanism 2. CA1 cells with complementary firing rate maps (top row) both significantly cross-coupled to the same mPFC cell (bottom row). Crosscorrelograms as in **J)**. Throughout the figure, ** = p<0.01, *** = p<0.001.

We considered an artificial agent that traversed two opposite 1D trajectories (for example, south to east and north to west in our task) in order to travel towards a goal. The spatial response probability of *selective* units was generated so as to resemble the firing of CA1 place cells, independently for each trajectory. This procedure created CA1-like units with very different symmetry properties, ranging from completely asymmetric to almost symmetric. The internal connectivity among *abstract* units was selected randomly and kept fixed during each synthetic experiment. The transfer of spatial information from *selective* towards *abstract* units was controlled by a matrix **C** (Fig. 3A). Our aim was to find the best possible **C** which maximized the mutual information between the spiking of *abstract* units and the distance to the goal, independently of the trajectory. We randomly initialized the spatial- and trajectory-selectivity of *selective* cells and the internal connectivity of the *abstract* cells 100 times, and optimized the mutual information by finding the best **C** via a sequential least squares programming routine (35).

First, we asked whether those optimal **C**s could recapitulate the observations made on the data. Indeed, we observed that couplings were strongest preferentially starting from *selective* units with symmetric firing fields (Fig. 3B), and that *abstract* units with strong couplings showed a higher single cell spatial information score (Fig. 3C). We further used the model to generate predictions to then be tested on the data. If the spatial properties of mPFC cells are inherited from CA1, one should be able to (at least partially) recover the firing rate maps of mPFC cells from the firing rate maps of CA1 cells and their across-area couplings (Methods). We found this to be the case, both in the model (Fig. 3D) and in the data (Fig. 3G), with a prediction that greatly outperformed random couplings. Further, we asked how the coupling to non-symmetric cells might be structured. For a mPFC cell to generalize its position-dependent firing across trajectories, it needs to receive information from two (or more) CA1 cells that are complementary to each other, i.e. fire in similar positions of opposite trajectories. To test this hypothesis, for each *abstract* unit we picked the two *selective* units with the strongest coupling. We measured the “complementariness” of these two *selective* units by measuring the similarity of their firing in opposite arms. We found them to be more complementary than chance (Fig. 3E), and observed the same pattern in real data (Fig. 3H). Accordingly, if two CA1 cells are complementary to each other, they should couple in a similar way to all the mPFC cells. To test this, we scatter-plotted the complementarity of each CA1 cell pair against the similarity of their couplings to all other mPFC cells. We found the two quantities to be significantly correlated, both in the model and in the data (Fig. 3F-I). Altogether, these results support the notion that the mechanisms by which a mPFC cell forms a generalized spatial code is by either coupling with a CA1 cell that by itself is more symmetric (Fig. 3J), or by coupling to more than one CA1 cell that together are complementary (Fig. 3K).

### CA1 - PFC coupling is task phase dependent

We explored the possibility that the coupling between the two areas is modulated by the task phase. We could confirm this by measuring the average excess of correlation among CA1-mPFC cell pairs in the start and goal arms separately: coupling was strongest and shortest in delay in the start and became weaker and more delayed in the goal arm, after the animal made its choice (start: ~25ms, goal: ~75ms) (Fig. 4A). Interestingly, we found that noise correlations detected separately in the two arms to be weakly (yet significantly) correlated when considering no delay (Fig. 4B), whereas they were not significantly correlated at their best delays (Fig. 4C), indicating a potentially different pathway of information transfer between the two areas during the two task phases.

**Fig. 4.**
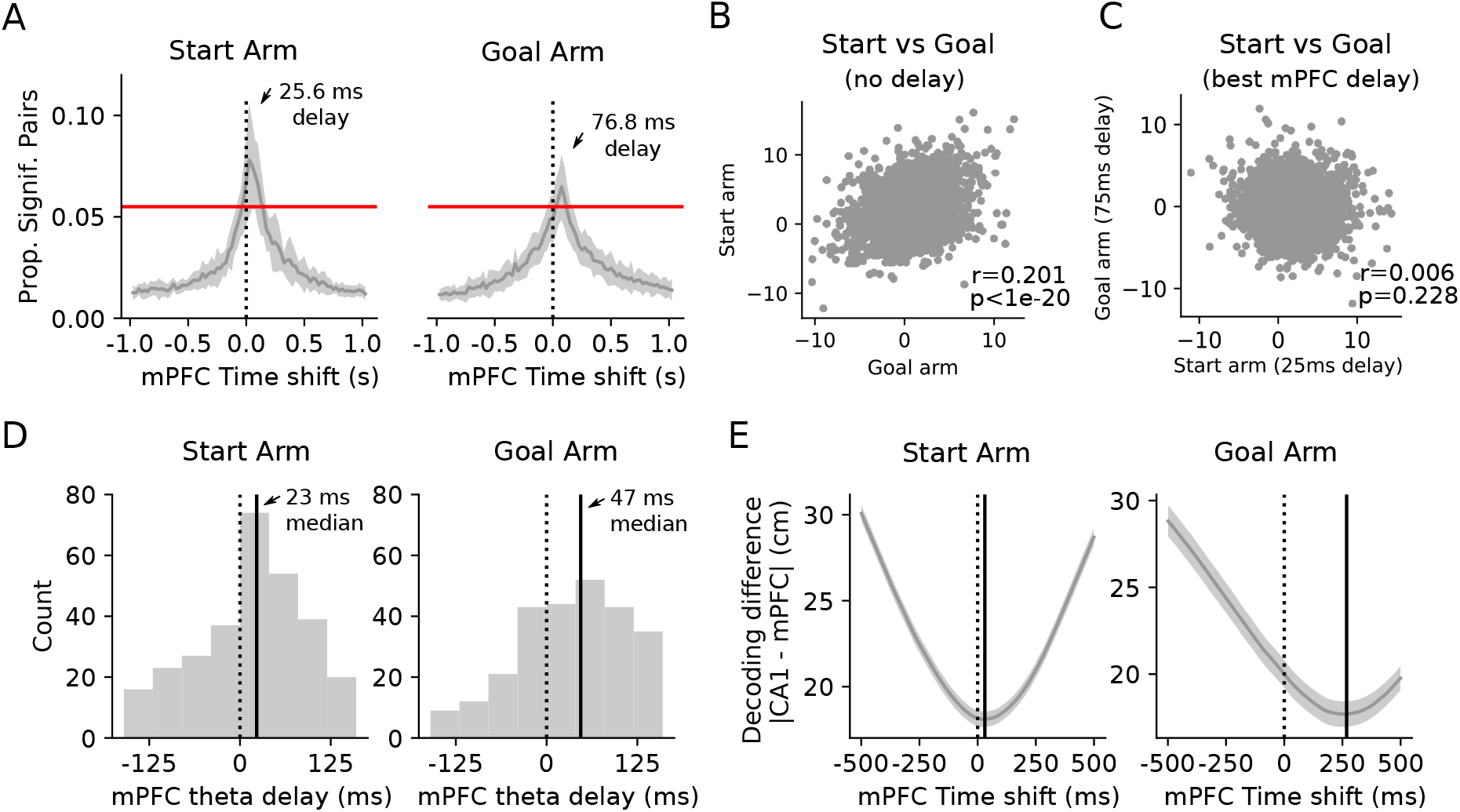
Task phase modulates cross-area couplings intensity and delay. **(A)** Proportion of significant cell pairs in start arm (left) vs goal arm (right) as a function of mPFC delay. A pair was deemed significant at a given delay *t* if the real data cross-correlation at *t* exceeded +3STD or fell below -3STD of the average cross-correlations at *t* computed on simulated data. Red line represents the average+5STD of significant couplings over all possible cell pairs and delays >500ms. Shaded area represents the 95th confidence interval for the mean proportion of cross-coupled cells computed across 13 sessions. **(B)** Scatter plot of noise correlations detected separately in the two arms without delay. Spearman correlation test, p<0.001, r=0.201. **(C)** Scatter plot of noise correlations detected separately in the two arms at their best mPFC delays (25.6ms for the start arm, and 76.8ms for the goal arm). Spearman correlation test, p>0.05, r=0.006. **(D)** Optimal locking of mPFC cells to the underlying CA1 theta oscillations. The delay that produced the best theta locking of each mPFC cell was found by computing a phase histogram for each delay and measuring the mean vector length (32). Mann–Whitney U test for start vs goal arm optimal theta delay, p<0.001. KS test for the distributions, p<0.001. **(E)** Alignment of CA1 - mPFC decoded position as a function of mPFC delay. The shaded area corresponds to the 95th percentile confidence interval of the mean over 13 sessions.

To further validate this observation, we employed a measure to detect the best theta locking of mPFC cells to the underlying CA1 theta oscillation (32). Theta coupling between the two areas is known to change in strength depending on the cognitive demand (36, 37). mPFC cells were preferentially locked with a median delay of ~23 ms in the start arm, whereas in the goal arm locking was weaker and more delayed (median ~47ms) (Fig. 4D). We also analysed the properties of the spatial information encoded by the two areas by measuring the alignment of decoded position as a function of mPFC delay separately for the start and goal arms. mPFC representation was more delayed and less coherent with CA1 in the goal arm as compared to the start arm (Fig. 4E). These observations corroborate the idea that both coupling and encoding coherence are stronger and shorter in delay before reaching the center of the maze, where the animal makes its upcoming trajectory decision.

### Spatial coding across rules

It has been shown that mPFC firing changes during rule-switching tasks (38–44), suggesting that this area encodes information about the rule or the insight that the rule has changed. Conversely, the hippocampus is well-known for its ability of changing firing characteristics in response to environmental and contextual changes (45–47). Here we tested the influence of the introduction of a new rule on the encoding of space by CA1 and mPFC populations (Fig. 5A). We computed the average firing in each location for all cells and compared the similarity of population vector activities within rules versus across rules (48, 49). We observed that CA1 population firing was significantly different across the two rules (Fig. 5B), showing rule-related population remapping in the hippocampus, whereas mPFC population vectors did not change more across rules than within (Fig. 5C). This effect might allow the mPFC to generalize across rules, providing a stable representation of the distance to the goal irrespective of the rule. To test this hypothesis, we decoded the distance to the goal from the activity of the two areas either using a decoder trained on different trials of the same rule or on trials of the other rule (Fig. 5A).

**Fig. 5.**
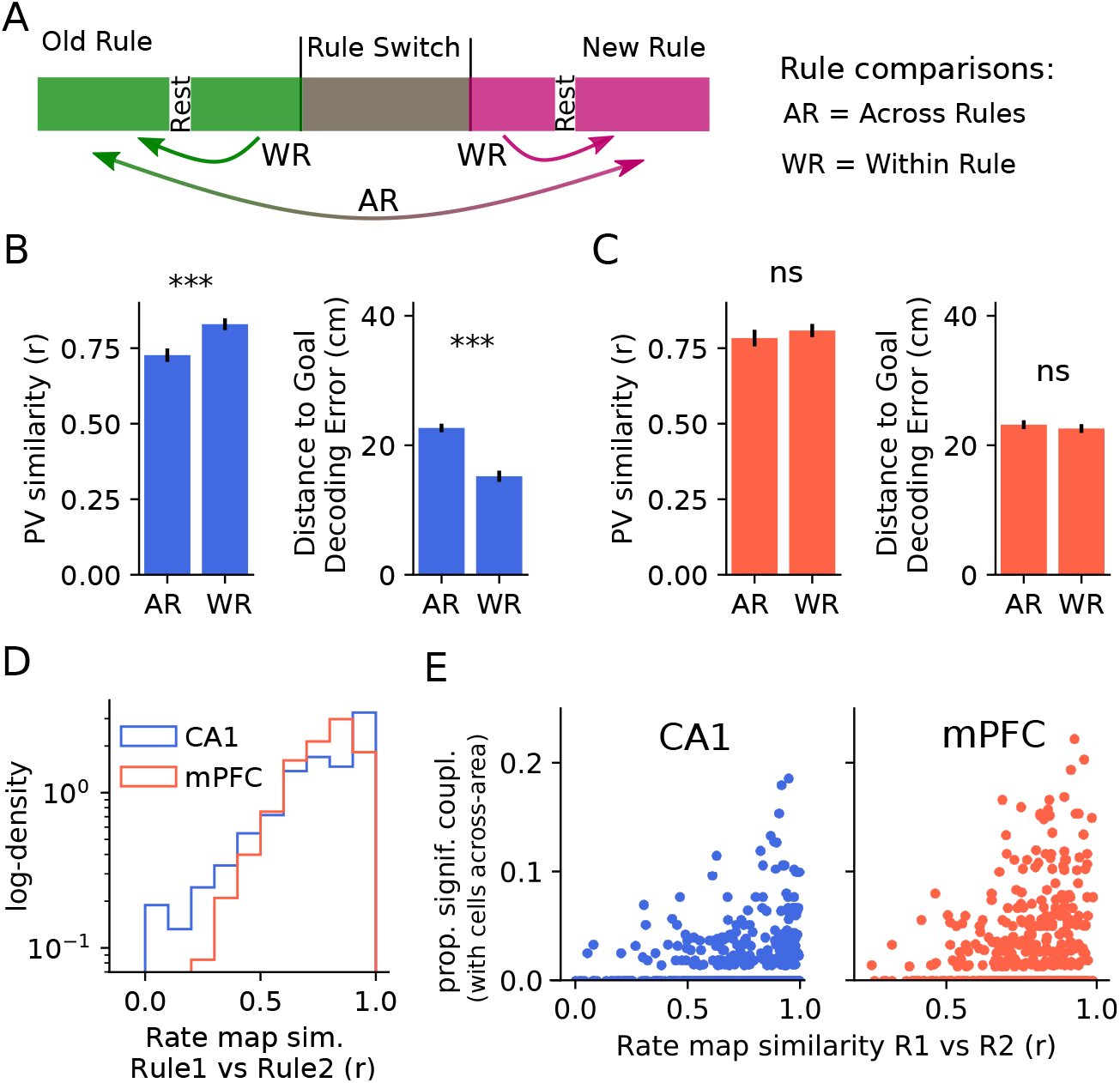
Spatial representation across rules. **(A)** Schematic of rule comparisons. Trials belonging to one rule were compared to different trials belonging to the same rule (WR) or different rule (AR). **(B)** Left: average similarity of CA1 population activity vectors (PVs) across rules (AR) versus within rules (WR). Error bars represent 95th confidence interval of the mean over PVs across locations and sessions. Mann–Whitney U test, p<0.001. Right: Average decoding error of a distance-to-goal bayesian decoder using CA1 cells activity (not used for training) from trials of the same rule (WR) or the other rule (AR). Error bars represent 95th confidence interval of the mean over PVs across locations and sessions. Mann–Whitney U test, p<0.001. **(C)** Same as B) but using mPFC data. Both comparisons tested using a Mann–Whitney U test, p>0.05. **(D)** Distribution of rate map similarity across rules of single pyramidal cells in CA1 (blue) and mPFC (red). KS test, p<0.001. **(E)** Rate map similarity across rules for each pyramidal cell against proportion of significant couplings with cells across-area (left: CA1, right: mPFC). Spearman’s rank correlation, CA1: *ρ* = 0.114, p<0.01; mPFC: *ρ* = 0.239, p<0.001. Throughout the figure, ns = not significant, *** = p<0.001.

We found that decoding performance from mPFC data remained stable (Fig. 5C), whereas it decreased in precision when doing the same on CA1 data (Fig. 5B). To deepen our understanding of remapping dynamics across rules, we compared the distribution of rate-map similarity of CA1 and mPFC single cells before and after rule switch. We found the distribution for CA1 to have a thicker left tail, explaining the difference in PV similarity across rules. Nonetheless, some mPFC cells also showed different spatial-firing dynamics across rules (Fig. 5D), which could represent rule or other contextual information. We characterized the relationship between cross-area coupling strength and single-cell remapping. For each putative pyramidal cell, the proportion of significant connections with cells in the other area was plotted against the rate map similarity between the old and new rule (Fig. 5E). We found that cells with the strongest coupling to the other area tended to remap less, both in CA1 and mPFC. This suggests that cross-area couplings are not influenced by changes in the current rule. To corroborate this hypothesis, we tested whether the underlying structure of CA1-mPFC coupling changes across rules. We computed the noise correlations separately for each rule and found the two values to be strongly correlated (Supp. Fig. 3A). The correlations remained strong also when we calculated them separately in the start and goal arms (Supp. Fig. 3B, C).

## Discussion

### Origins of the prefrontal generalized code

Here we showed multiple pieces of evidence supporting the long standing hypothesis that the hippocampus provides the prefrontal cortex with spatial information (19, 21).

First, we found that the decoding errors in the hippocampus and prefrontal cortex were correlated, and this correlation improved when mPFC responses were delayed by ~ 125ms. Moreover, the spatial coding in the prefrontal cortex lagged behind that of the hippocampus by ~ 100ms (Fig. 1). These results confirm and expand upon previous studies (21). These results were reinforced by transfer entropy measures, which indicated that most of the information flow was directed from hippocampal to prefrontal cortical neurons (28, 50).

Second, we found robust statistical evidence for the existence of pairwise, across-area coupling between neurons. Previously, mPFC-hippocampus coupling has been investigated either by lesioning or inactivating projections or brain regions (11, 18, 22, 23, 51), or by measuring theta coherence or selectivity (32, 36, 37). Here we used noise correlation analysis to find hippocampal-prefrontal cell pairs that were functionally coupled, i.e., cells whose pairwise correlation was much higher or much lower than expected from our null model simulations (27, 52). Intriguingly, mPFC cells that were significantly cross-coupled to the hippocampus held more spatial information (Fig. 2). The majority of couplings were found at a mPFC delay of 25~75ms, likely attributed to a multi-synaptic transmission of information. Moreover, mPFC cells showed a positive correlation between theta locking and coupling strength, making theta locking a candidate mechanism underlying mPFC and hippocampal cells coupling (32). The fact that the spatial representation was slightly more delayed than cellular couplings might be due to internal neocortical computations (53), similarly to what has been observed in entorhinal cortices (54). It remains to be tested whether cells in mPFC that are cross-coupled are the same that show enhanced coordination with hippocampal sharp-wave ripples (24, 55).

Third, the structure of across-areas couplings, and the properties of the cells involved were predicted by an optimal model of information transfer and abstraction (Fig. 3). This modeling approach allowed us to investigate the potential mechanisms behind generalization. We found that generalization was achieved in prefrontal units either by connecting more strongly to cells in the hippocampus that showed symmetric and trajectory-independent firing; or, alternatively, by connecting to pairs of hippocampal cells whose firing across arms was complementary. The suggested processes might be at the core of generalization procedures across hippocampal and neocortical areas (16, 56, 57), which will need to be confirmed in future studies.

Whilst the prefrontal cortex might inherit spatial information from the hippocampus, these findings by no means exclude the fact that other types of spatial and non-spatial codes travel from the prefrontal cortex to the hippocampus. There are several indirect connections between the prefrontal cortex and the hippocampus, enabling bidirectional information flow (14). For example, it is known that inactivating the medial prefrontal cortex or the indirect nucleus reuniens projections leads to a decrease in hippocampal place cell firing variability (9, 51), rule-based object selectivity (23) and trajectory-dependent firing (11).

### Different routing of information upon changes in cognitive demands

In the present task, animals need to reach the center and decide where to turn. It is reasonable to assume that cognitive demand is higher upon reaching this decision point (36). We observed that the functional connectivity between the hippocampus and prefrontal cortex was strongest and less delayed in the start arm compared to the goal arm (Fig. 4). This may indicate that hippocampal-prefrontal communication is enhanced before a spatial decision is made and when cognitive demand is high. This is in line with studies on theta oscillatory synchronization that showed enhanced coupling before the choice point or when rules have changed (36, 37). It is also possible that the routing of spatial information follows different paths with different types of cognitive load. In support of this observation is the fact that direct projections from the ventral hippocampus to prefrontal certex are important only for task periods requiring memory encoding (18). From an anatomical point of view, there are several possible routes via which the hippocampus could transmit spatial information to the mPFC. Apart from the prominent direct projections arising from the ventral hippocampus (13), indirect projections can be established through lateral entorhinal and perirhinal cortices (13, 17). Information may also be transferred indirectly via the thalamic nucleus reuniens; this area bidirectionally connects with the mPFC (15, 58), projects to the hippocampus (15) and in turn receives hippocampal input mostly via the subiculum (14). Whilst there is evidence that information in the mPFC passing to the hippocampus via the reuniens is required for certain aspects of hippocampal spatial firing (11), there has been no demonstration yet for a flow of spatial information via this route from the hippocampus to mPFC.

### Novel insights on hippocampal and prefrontal spatial coding

Across area couplings preferentially involved hippocampal cells that showed more generalized spatial coding. This result underlies our finding that in the present task a subset of hippocampal cells generalize spatial location, i.e., fire similarly in the same position of different arms, in agreement with (56, 59). This finding resembles and sheds light on the similarity of firing across same-direction runs of certain hippocampal cells on the hairpin maze (60). Moreover, we also found a small yet significant subset of hippocampal cells that encoded egocentric position (Fig. 1). This is different from previously reported splitter cells (47, 61, 62), in that here we found that certain cells had the same encoding for trajectories that require the same turn, instead of differentiating all the trajectories. Our finding is reminiscent of other studies of hippocampal cells with egocentric coding in mice (63), and others in bats, where a subset of neurons showed egocentric goal direction selectivity (64). Altogether, this corroborates the idea that hippocampal activity is also important for egocentric coding (65). We also expanded upon our understanding of the generalization of mPFC spatial representation (24–26), and its extent, by comparing the trajectory-specificity of the maps and their effect on the decoding (Fig. 1). Studies in monkeys, using novel measures of the geometry of responses, showed that not only prefrontal areas like the dorsolateral mPFC and anterior cingulate cortex (ACC), but also the hippocampus hold a generalized representation of task-relevant variables (57). That the hippocampus in the present task is less prone to generalization might underlie specific task requirements, or the fact that spatial information requires more processing steps to reach that level of abstraction.

### Stability of couplings across rules

It has previously been shown that the hippocampus remaps upon changes in the environment (45, 46, 66). We observed that the hippocampus remaps with changes in the rule, despite an unchanging environment. This is similar to (26), and is reminiscent of previous works, where remapping occured after learning in spite of environmental and contextual cues remaining the same (49, 67). The mPFC, at a population level, did not show significant changes in spatial firing propensity for different rules, in apparent contrast with other studies (38–44). This discrepancy might be explained by the fact that in some of these studies the behavioural tasks were different. Moreover, we only considered the representation of well-established rules, which does not exclude the possibility that the coding differs during shorter periods during rule switching. Furthermore, analysing single cells we found some differences in the average spatial firing of prefrontal cells across rules. A potential explanation for the fact that in this task the average population spatial representation changes less in mPFC across rules might be that, intrinsically, the representation is in an abstract form, allowing the mPFC to faithfully decode the position of the animal in spite of contextual changes. We found that to be the case, with a negligible loss of spatial decoding precision (Fig. 5). For the hippocampus, on the other hand, the contextual precision of the spatial representation came with a cost: decoding the position in one rule with a decoder trained on the other rule resulted in a significant loss of precision. Nonetheless, although hippocampal cells showed rule-related remapping of their spatial firing properties, the connectivity between hippocampal and prefrontal cells was more stable than chance and both hippocampal and prefrontal cells with stronger cross-couplings remapped less. This suggests that CA1 cells participating in cross-area functional coupling have a more stable spatial code, thus providing reliable spatial information for the mPFC. Accordingly, our results also show that cross-coupled spatially-selective mPFC cells are less sensitive to changes in the rule,allowing for a context-independent and generalized representation of space.

## ACKNOWLEDGEMENTS

We thank Federico Stella for invaluable suggestions and discussions. We thank Yosman BapatDhar and Andrea Cumpelik for comments, help and suggestions on the exposure of the text. We thank Predrag Živadinović and Juliana Couras for comments on the text and the figures. This work was supported by the EU-FP7 MC-ITN IN-SENS (grant 607616).

## Materials and Methods

### Experimental methods

The data used in this study is the same as used in (25). We will report the experimental methods here for completeness.

#### Subjects and Surgery

Four male Long-Evans rats (300-350 g, 2-4 months of age; Janvier, France) were used in this study. The animals were housed in a separate room on a 12 hour light/dark cycle and were taken to the recording room each day prior to the experiments. Animals shared a cage with littermates before surgery. All procedures involving experimental animals were carried out in accordance with Austrian animal law (Austrian federal law for experiments with live animals) under a project license approved by the Austrian Federal Science Ministry (License number: BMWFW-66.018/0015-WF/V3b/2014). Rats were implanted with microdrives housing 32 individually-movable tetrodes, arranged into three bundles targeting the right dorsal hippocampus (specifically dorsal CA1, HPC) and left and right medial prefrontal cortex (specifically prelimbic area, mPFC). The HPC bundle consisted of 16 tetrodes and the two mPFC bundles of 8 tetrodes each. Tetrodes were fabricated out of four 12 mm tungsten wires (California Fine Wire Company, Grover Beach, CA) that were twisted and then heated to bind into a single bundle. Tetrode bundle lengths were cut so that the two mPFC bundles were 1-1.5 mm longer than the HPC bundle. The tips of the tetrodes were gold-plated to reduce the impedance to around 300 kU. Before surgery the animal was put under deep anesthesia using isoflurane (0.5%–3%), oxygen (1–2 L/min), and an initial injection of buprenorphine (0.1 mg/kg) and ketamine/xylazine (7:3 ketamine (10%) and xylazine (2%), 0.05ml/100 g). Craniotomies were drilled above the HPC (AP: 2.50 to 4.50, ML: 1.2 to 3.6) and above the mPFC across the sinus (AP: 4.60 to 2.50, ML: 0 to ± 0.8). Six anchoring screws were fixed onto the skull and two ground screws were positioned above the cerebellum. After dura removal the tetrode bundles were centered above their respective craniotomies and lowered into the brain at a depth of 2 mm for the mPFC and 1 mm for the HPC. The exact depth of mPFC tetrode implantation was noted to ensure later lowering into the target area. Tetrodes and craniotomies were coated in paraffin wax and the microdrive was anchored to the skull and screws with dental cement. The analgesic meloxicam (5 mg/kg) was given up to three days after surgery and the animal was allowed one week recovery. Thereafter, tetrodes were gradually moved in 50-200 mm steps into the HPC pyramidal cell layer and mPFC.

#### Plus maze apparatus and task

Following the recovery period, animals were food-restricted with ad libitum access to water and accustomed to the plus maze and rest box. The plus maze was elevated (80 cm) and consisted of four arms (85 cm long and 12 cm wide), referred to as north, east, south and west, and a connecting center. The animal was placed in one of the two start arms (north or south) and had to collect a food reward (MLab rodent tablet 20mg, TestDiet, Richmod, USA) in one of the two goal arms (east or west), depending on the rule employed. Access to the arm not chosen as the start was restricted, so that the maze became T-shaped. A small light at the end of one of the two goal arms was switched on. Which arm was chosen as the start and light-on arm was chosen pseudorandomly for every trial, ensuring that an arm was not chosen more than three consecutive times. Once the animal reached a goal arm and 5s passed, the animal was manually picked up and placed in the rest box before commencing to the next trial after a delay of 10s. The animal had to retrieve the reward based on a spatial or response (light) rule. During the spatial rule the reward was always placed in either the east or west arm, while during the response rule the reward was placed in the light-on arm. Importantly, also during the spatial rule one of the two arms was lit, but did not necessarily indicate the location of reward. To prevent an odor-guided strategy pellet dust was scattered along the maze and pellet-filled cups invisible to the animal placed under both goal arms. On each recording day, the animal underwent behavioral blocks as follows: rest, rule 1 (previous day’s old rule), rest, pre-switch, rule switching, post-switch, rest, rule 2 (new rule), rest. After the first rest, the animal started by performing trials based on the previous day’s old rule. After reaching performance criterion (see below), the animal rested again and afterwards the rule switching phase began. During the pre-switch block the animal had to collect reward based on the last rule of the previous day until reaching the performance criterion (see below). Then the rule was changed and reward had to be collected based on the new rule. The change in rule was not announced to the animal, which had to switch to the new rule through trial-and-error until performing to criterion. Trials performed after the rule change, but before the animal reached good performance comprised the rule switching block, while the post-switch block comprised all trials from the beginning of good performance ( defined in the next sub-subsection*). The animal had to perform cross-modal switches, i.e., switches from spatial to light or light to spatial rule, never between the two spatial rules. While correct performance of a spatial rule involves two trajectories (e.g., go-east rule: north to east and south to east), correct performance of the light rule can involve any of the four trajectories. Therefore, the performance criterion for the spatial rule was set to 12/15 and for the light rule to 24/30 correct trials, ensuring a similar number of light rule trials where the animal performed trajectories that matched those of the spatial rule. After another rest session, the animal performed a final 20 trials of the newly acquired rule.

### Beginning of good performance

All trials before the rule change comprised the pre-switch block. Trials performed after the rule change, but before the animal reached good performance comprised the switching block. The beginning of good performance (*bgp*) was defined as the center index after rule change where the error rate over five consecutive trials dropped to zero.

#### Histology and reconstruction of recording positions

After the final recording day tetrodes were not moved. Animals were administered ketamine/xylazine (7:3 ketamine (10%) and xylazine (2%), 0.1ml/100 g) and overdosed with pentobarbital (300mg/ml) before being transcardially perfused with 0.9% saline followed by 4% formaldehyde. Brains were extracted and stored in 4% formaldehyde. On the same day brains were transferred into 30% sucrose solution until sinking for cryoprotection. Finally, brains were quickly frozen, cut into coronal subsection*s with a cryostat (50-60 mm), mounted on glass slides and stained with cresyl violet. The positions of tetrode tips were determined from stained subsection*s and cells recorded from tetrodes outside mPFC were excluded from analysis. For cells recorded from HPC tetrodes the presence of SWRs in the field recordings served as inclusion criteria.

#### Data Acquisition

The extracellular electric signals from tetrodes were pre-amplified using a headstage (4 x 32 channels, Axona Ltd, St. Albans, Hertfordshire, UK). The amplified local field potential and multiple-unit activity were continuously digitized at 24 kHz using a 128-channel data acquisition system (Axona Ltd). Two red LED bundles mounted on the preamplifier head-stage were used to track the *x,y* location of the animal. Every day before recording, HPC tetrodes were moved optimizing the yield of recorded cells. Additionally, mPFC tetrodes were lowered every day by 30-50 mm to ensure recording of a new population of cells.

#### Spike sorting and unit classification

Clustering of spikes and unit isolation procedures were described previously (68). Briefly, the raw data was resampled to 20 kHz and the power in the 800-9000 Hz range was computed for sliding windows (12.8 ms). Action potentials with a power of > 5 standard deviations (SD) from the baseline mean were selected and their spike features extracted with principal components analysis. Action potentials were then grouped into multiple putative units based on their spike features using an automatic clustering software (http://klustakwik.sourceforge.net; (69)). The generated clusters were then manually refined using a graphical cluster-cutting program and only units with clear refractory periods in their autocorrelation, well-defined cluster boundaries and stability over time were used for further analysis. An isolation distance (based on Mahalanobis distance) was calculated to ensure that spike clusters did not overlap (69). Putative excitatory pyramidal cells and inhibitory interneurons were discriminated using their auto-correlograms, firing rates and waveforms (70). In our analysis we included only cells with an average firing rate > 0.25spikes per second in each of the three experimental phases. This comprised a total of 530 hippocampal (putative) pyramidal cells and 160 interneurons, and 477 prefrontal (putative) pyramidal cells and 105 interneurons.

### Statistical analysis

#### Linearized Position

To linearize the behavior of the animal, we calculated the distance from the center from the 2D spatial position of the animal (25). This way a “V-shaped” positive function for each trial was obtained. For each position before the center (i.e., before the global minimum) we subtracted the minimum and then changed the sign. Then, 100 was added to every position to obtain a positive measure of the relative position of the animal between start (0 cm) and goal (200 cm). The center corresponded to 100 cm.

#### Firing rate maps

We inferred the average firing of each cell at each given location of the environment separately for three cases: 2D maps, 1D trajectory dependent and 1D trajectory independent. In every case, we utilized data from periods when the animal was moving faster than 7cm/s to avoid potential nonlocal population activity (25).

##### 2D maps

We binned the *x, y* locations of the animal in 5cm square bins. We counted how much time the animal spent in each location, which corresponded to the occupancy map. Afterwards, for each cell, we counted the number of spikes emitted in each location, and divided by the time spent there. Finally, we regularized by convolving with a 2D Gaussian kernel with *σ* = 2bins.

##### Trajectory-dependent 1D maps

Since the arms of the maze are relatively narrow, we computed a linearized version of the firing rate maps, effectively yielding an average firing as a function of the distance to the goal, separately for the 4 trajectories: North to East(NE), North to West (NW), South to East (SE), South to West (SW). We selected the trials where the animal followed one trajectory, and with that data binned the linearized position in 10 cm bins and counted the time spent at each discrete location. Afterwards, for each cell and each trajectory, we counted the number of spikes emitted in each binned location and divided by the time spent there. Finally, we convolved the rate maps of each cell for each trajectory with a Gaussian kernel with *σ* = 1bin.

##### Trajectory independent 1D maps

We computed the same quantity as above, without separating the trials into 4 trajectory-groups.

##### Symmetry of firing rate maps

We measured the “mirrordness” of firing rates to check whether single cells fire similarly at opposite sides of the track. To do that, we computed the Pearson correlation of the linearized firing rate maps of opposite arms (S vs N and E vs W) and reported the average of the two measures.

##### Allocentric vs Egocentric maps

To test whether cells fired as a function of the turning direction, instead of the absolute location on the maze, we analyzed the trajectory-dependent linearized firing rates. In particular, we reasoned that a cell with egocentric coding would fire similarly on the two different goal arms during trials with the same turn, and would fire differently on the same arm for trials with different turns. On the other hand, if a cell had an allocentric coding, it should always fire similarly on the same goal arm independently of turning direction. To formalize our intuition, we considered the trajectory dependent rate maps and, for each cell, we measured the Pearson correlation of the position-dependent firing on the same goal arm (110 – 180cm) for different turns and subtracted the correlation of opposite goal arms for the same turn. This way we obtained a measure within –2 and 2, which was negative for egocentric cells and positive for allocentric cells. If a cell showed symmetric behavior (i.e. same firing on opposite arms, independent of turn) this measure would be close to zero. To test how many cells were significantly egocentric or allocentric, we compared the measure against the distribution obtained by shuffling the trial identities 1000 times while keeping everything else fixed. We labelled cells as significantly allocentric if their measure was higher than the 95th pecentile of their own distribution, and significantly egocentric if their measure was lower than the 5*th* percentile. Finally, we tested for significance by employing a Binomial test with *p* = 0.05 on the number of cells that were significantly allocentric (or egocentric) out of the total number.

##### Complementarity measure

To measure if two cells are “complementary”, i.e., if their joint activity could help to form a symmetric pattern, we computed the Pearson correlation of the linearized firing rate maps for opposite trajectories (for example, SW for cell 1 vs NE for cell 2) and then reported the average of the 4 possible comparisons.

##### Spatial information measure

We computed the spatial information per spike as described in (71, 72). In a nuthshell, this corresponds to the first order approximation of the mutual information between position-dependent spiking probability and location, divided by the average firing rate. Denoting with *s* the location, and *λ_s_* the average firing of a cell at location *s*, and *λ* its average firing rate, the spatial information is defined as

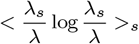

where < · >_*s*_ denotes an average over positions.

#### Decoding of distance to the goal

For each decoding scheme we randomly selected 75% of the trials for computing the firing rate maps, and used the remaining 25% for decoding and assessing decoding quality. We employed a maximum a posteriori (MAP) Bayesian decoding under the assumption that cells are independent and fire according to a Poisson distribution (73).

#### Correlation of decoding errors

To check whether the errors of one area agree with the errors of the other area, and whether this effect increases at a delay, we employed a delayed version of an analysis proposed by Zielinski and colleagues (21). If the spatial encoding of PFC is delayed compared to CA1, in that either PFC receives information from CA1, or that information takes longer to reach PFC, then spikes in PFC should represent spatial information that is older than CA1’s. Hence, we took all PFC spiking times and added a time lag *τ* ∈ [–500, 500]ms, and for each lag we computed firing rate maps on a fraction of data (random 75%) and decoded the position from the remaining part of the data. Afterwards, for each delay we computed the Spearman correlation between decoding errors from CA1 (without lag) and the lagged PFC activity. Confidence interval correspond to the 95th percentile for the mean taken over the 13 sessions analyzed.

#### Detection of cross-area couplings

We employed a statistical modelling approach to detect pairs of CA1-PFC cells that are significantly coupled. With “coupled” we denote cell pairs that are significant noise correlated (29). For each cell we fitted a generalized linear model (GLM) (30, 31) that included all possible covariates measured which could influence and explain the cross-area correlations. These covariates were: linearized spatial position, trajectory, theta selectivity, speed selectivity, spiking history and within-area spiking of other cells (i.e. PFC cells were fitted with the spiking of other PFC cells only and, separately, CA1 cells with the spiking of the other CA1 cells only; for further details see subsection “Modelling” below).

These models were used to compute a rigorous statistical test, the null hypothesis being that the cross-correlogram among cell pairs is completely explained by external covariates. The alternative hypothesis is that external covariates cannot explain the amount of covariability, hence we considered that a signature of “coupling”. This approach is similar to the one introduced in (27).

With these GLM null models, we simulated the activity of each cell 10000 times and, for each CA1-PFC cell pair, a cross-correlogram of the responses was computed. Those surrogate cross-correlograms were used to measure by how much the actual pairwise cross-correlation measured on real data differed from the simulated ones. We did this for each possible PFC delay in the range of ±1sec.

We considered a cell pair significantly coupled if the peak of the actual cross-correlogram exceeded the mean plus 4 standard deviations of the peaks of the 10000 surrogate cross-correlograms for that pair. Cells in one area that showed significant noise correlation to at least one cell in the other area were termed “cross-coupled”.

#### Theta locking and optimal delay

We employed a measure introduced by Siapas and colleagues in (32). Briefly, we filtered the hippocampal LFP between 5 and 15 Hz and computed the phase of each spike of each cell in PFC using a Hilbert transform. Afterwards, we employed a Rayleigh test for circular uniformity and selected all cells that yielded a *p* < 0.05. For each one of those cells, we introduced a delay *τ* ∈ {−125, −124,…, 124, 125}*ms* to all spikes, again computed the phase with a Hilbert transform and computed the mean vector length (MVL) of these angles. We then selected the delay that yielded the highest MVL for each cell, and computed a histogram, which we reported in fig. 4.

#### Population vector similarity

Population vector (PV) similarity is a measure that allows us to quantify the change in population activity at any given location across contexts. This measure has been employed to quantify remapping in a number of studies (48, 49). The PV at a particular location represents the vector of the average activities of all the cells in the population under study. We constructed them starting from the previously computed firing rate maps separately for the two populations and the two rules. We then employed a Pearson correlation to quantify the similarity of the PV for each location across ruless, and reported the mean ± 95th confidence interval for the mean.

#### Transfer Entropy

Transfer entropy is a non-parametric measure of directed (time-asymmetric) transfer of information between two random processes (28). Transfer entropy from a process *X* to another process *Y* is the amount of uncertainty reduced in future values of *Y* by knowing the past values of *X*, conditioned on past values of *Y*. More specifically, if *X_t_* and *Y_t_* denote two random processes, transfer entropy from *X_t_* to *Y_t_* is defined as the conditional mutual information between *Y_t_* and the history of *X_t_,* denoted by ***X***_*t*–1,*t*–2,…_, conditioned on the history of the influenced variable ***Y***_*t*–1,*t*–2,…_:

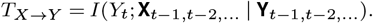

We used the JIDT package (50) to estimate this quantity, which was measured across binned (25.6ms) and binarized spike trains of each pair of CA1-PFC cells. Significancy was measured by comparing the actual value against the 1000 obtained by randomly shifting the spike trains (uniform random from 1 to 100 seconds)(49).

### Modelling

#### GLM model of cells response

We utilized a GLM model to describe each cell’s response propensity as a function of all measured covariates during foraging activity. These detailed models served as null models for the statistical test we employed to detect couplings among cells (see section “Detection of across-area couplings”). We are going to describe in detail here the covariates used to fit the model and the parameters used in fitting and simulation routines, and refer the reader to other references for the details regarding the theoretical background of GLMs (31). The model described the inhomogeneus Poisson activation rate *λ_t_* of cells in 25.6ms time windows, *τ* = 0.0256s,

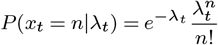

The expected firing rate *λ_t_* takes form

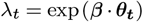

where ***β*** represents the model coefficients, found by maximum likelihood (below) and ***θ***_*t*_ represent the covariates, which are:

- trajectory-dependent spatial position: we binned the linearized position in 10cm bins, as described above, in 25.6ms time bins. We allowed each cell to have a different encoding for each trajectory separately to allow for maximum flexibility: this resulted in a 80-dim vector, i.e., 20 location bins for each possible trajectory. At each time point, only the entry of this 80-dim vector that represented the location of the animal, and the trajectory taken, was set to 1, and all the others to 0 (one-hot encoding variable)
- speed: we binned the speed, which was measured from the behavioral recording in 25.6ms time bins, in 7 non-overlapping and equally populated speed bins, starting from 7cm/s (one hot variable)
- theta phase: we computed the theta phase at the center of each 25.6ms time bin. The computation of the theta phase is based on Hilbert transform and is detailed in a previous subsection (“Theta locking and optimal delay”). We binned the angles in 10 nonoverlapping angular bins, and encoded it as a one-hot variable.
- spiking history: the spiking of the last three time bins was used
- whithin-area spiking activity: the spiking activity of all the other cells in the same area, together with the spiking history of each of those cells in the previous time bins were used as additional covariates

The number of parameters of such models ranged from 100 to 400, depending on the number of cells recorded simultaneously in the same area. We utilized the routine *GLM* offered by the package *statsmodels* v0.12.2 in Python 3.7 (74). We fitted the models by using an *L*2 regularization, whose aparameter was found by grid search on (10^-10^, 10^-9^,…, 10^0^) and cross-validation (train=75%, test=25% of data) independently for each cell.

#### Normative model of (spatial) information transfer and generalization

Consider two populations of neurons, exemplifying *N* PFC cells and *M* hippocampal cells. We will denote with 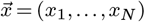 the stochastic (binary) activation of PFC cells, and with 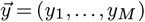 the stochastic (binary) activity of CA1 cells. Let us denote with *s* the distance to the goal, and with *k* the trajectory. We consider *s* to be a random variable that take values in {0,1,…,10}, each with equal probability, and *k* a Bernoulli(*p* = 0.5) ∈ {0,1}, independent of *s*.

We assume that 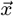 is driven by hippocampal input and internal connectivity, but has no initial selectivity for location. We formalize this request with a stochastic model, which is similar in its formalization to a restricted Boltzmann machine (34):

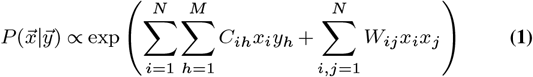

where *C* ∈ ℝ^*N×M*^ and *W* ∈ ℝ^*N*×*N*^.

We assume that the activity of CA1 cells is both trajectory and position dependent. We have that

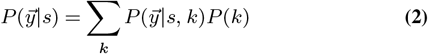

We also have that PFC population activity is position dependent through CA1, i.e. :

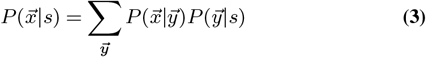

For fixed *W* and 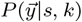, we want to find the best *C* that maximizes the mutual information between 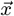 and *s*.

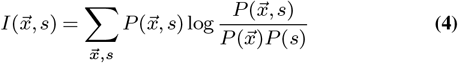

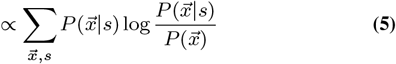

We maximize this quantity via Sequential Least SQuares Programming (SLSQP) routine in SciPy (75). We constrain each *C_ij_* to lay in [-1,1].

Our simulations use *N,M* = 10 neurons, which allows the mutual information to be computed without the need for approximations (by enumerating all possible patterns). Reported estimates are obtained by averaging across 100 randomly initialized networks; for each simulation, *W* is a symmetric matrix with entries randomly samples from a *N*(0, 1) distribution, and 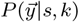 was initialized by considering each cell independent of each other, with a gaussian place fields per trajectory and simulated in such a way so as to resemble CA1 single cell statistics measured in the data.

## Supplementary figures

**Supp. Fig. 1.**
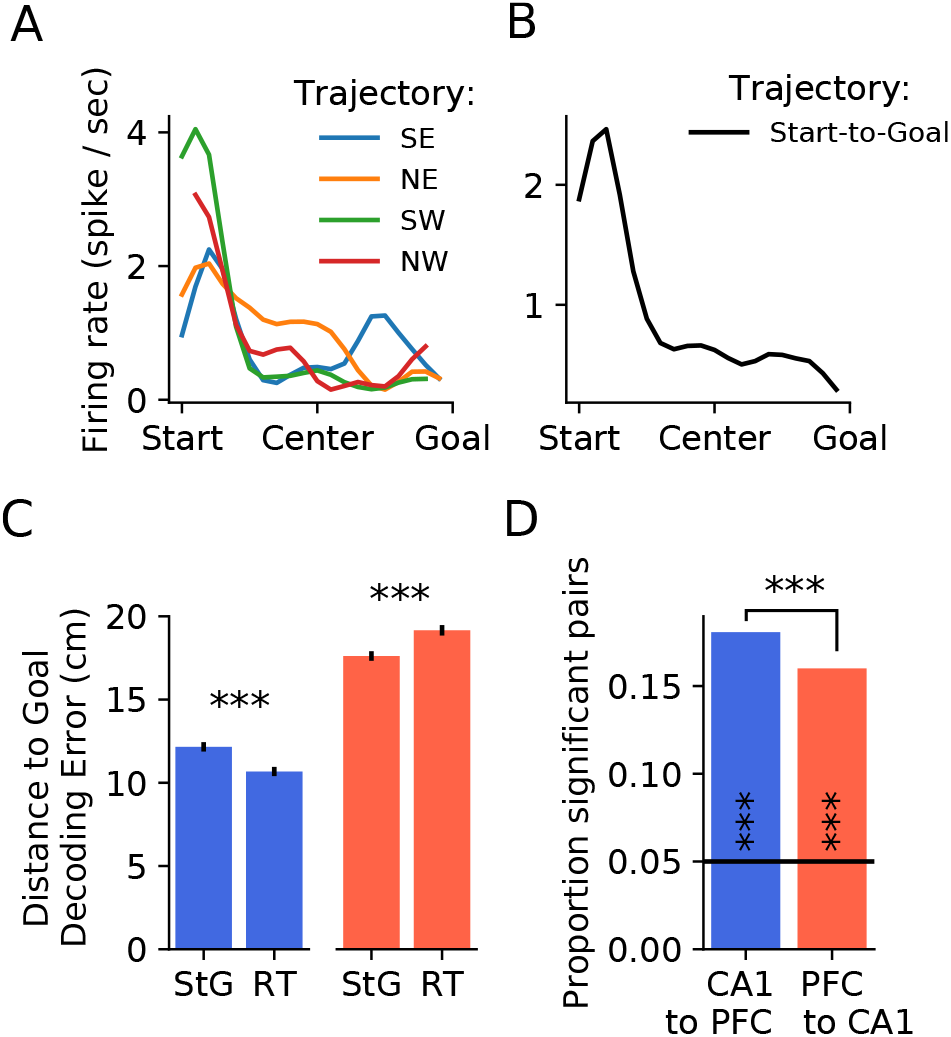
Further decoding schemes and Transfer Entropy measures. **(A)** Example mPFC cell’s firing rate as a function of distance to goal for each trajectory separately. **(B)** Firing rate as a function of distance to goal (independent of trajectory) for the same cell as in **A)**. **(C)** Average error of a bayesian decoder that took into account the trajectory identity (RT = Running Trajectory, same as **(A)**) or without information about trajectory identity (StG = Start-to-Goal, same as **(B)**). Both comparisons are significant under a Mann-Withney U test (both p<0.001). **(D)** Proportion of significant Transfer Entropy (TE) cell pairs across areas for the 2 directions. Black horizontal line denotes chance level. Each count was tested against chance using a Binomial(0.05) test (both p<0.001). The proportion of significant pairs CA1 to mPFC was significantly higher, as tested by using a Chi-Squared test for contingency tables (p<0.001).

**Supp. Fig. 2.**
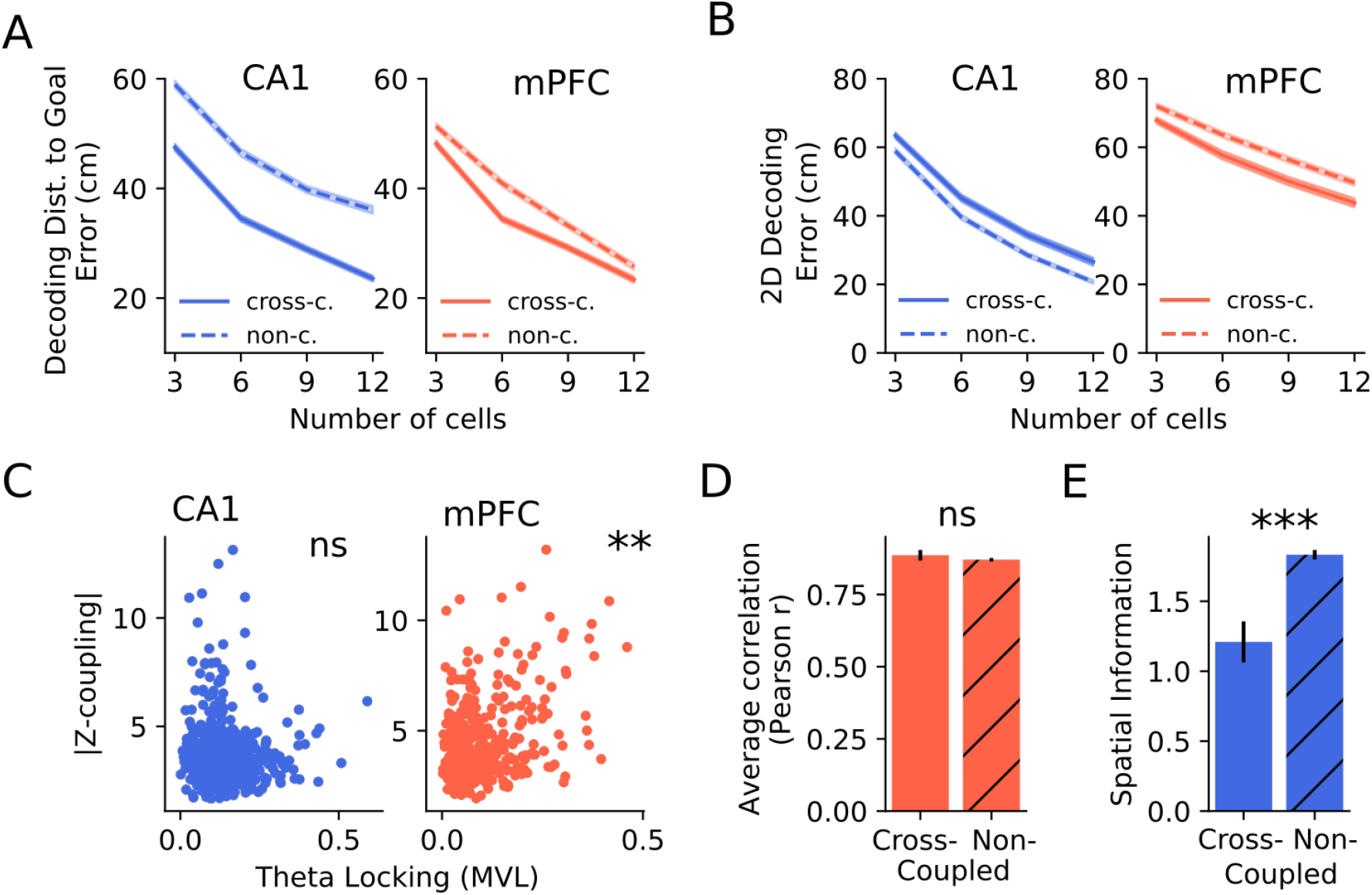
Decoding, theta selectivity and other properties of coupled cells. **(A)** average error when decoding the distance to goal with a StG strategy (see Supp. Fig. 1B), without considering the trajectory identity, for the same number of strongly vs weakly coupled cells, which were selected randomly 100 times among the subpopulation they belonged to. Left: CA1, right: mPFC. Solid line: cross-coupled cells; dashed line: non-coupled. Shaded area represents 95th confidence interval of the mean computed over 13 sessions. **(B)** same as **A)**, but decoding the (x,y) coordinate in space from 2D firing rate maps. **(C)** Phase locking to hippocampal theta of CA1 (left) and PFC (right) cells against the average absolute coupling to the other area. Pearson correlation test: CA1: r=0.015, p>0.05; mPFC: r = 0.271, p<0.01. **(D)** Symmetry of cross-coupled vs non-coupled PFC cells. Mann–Whitney U test, p>0.05. **(E)** 2D spatial information of cross-coupled vs non-coupled CA1 cells. Mann–Whitney U test p<0.001.

**Supp. Fig. 3.**
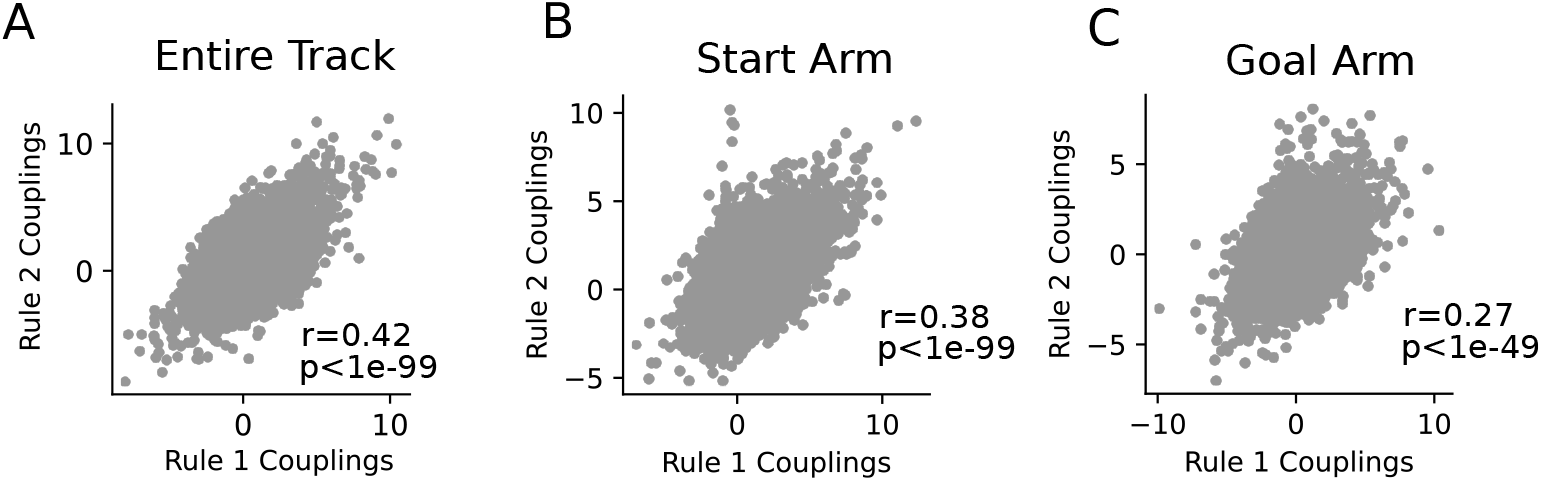
Stability of cross-area noise correlations. **(A)** Noise correlations (at 0-lag) measured on the entire track for every cross-area cell pair on rule 1 trials (x-axis) vs rule 2 trials (y-axis). **(B)** Same as A), computed on start arm only. **(C)** Same as A), computed on goal arm only. Figure panels include Pearson correlation coefficients and significance of Pearson correlation tests.

## Notes

### Competing Interest Statement

The authors have declared no competing interest.

